# Gray matter volume and estimated brain age gap are not associated with sleep-disordered breathing in subjects from the ADNI cohort

**DOI:** 10.1101/786707

**Authors:** Bahram Mohajer, Nooshin Abbasi, Esmaeil Mohammadi, Habibolah Khazaie, Ricardo S. Osorio, Ivana Rosenzweig, Claudia R. Eickhoff, Mojtaba Zarei, Masoud Tahmasian, Simon B. Eickhoff, for the Alzheimer’s Disease Neuroimaging Initiative

## Abstract

Alzheimer’s disease (AD) and sleep-disordered breathing (SDB) are prevalent conditions with rising burden. It is suggested that SDB may contribute to cognitive decline and advanced aging. Here, we assessed the link between self-reported SDB and gray matter volume in patients with AD, mild cognitive impairment (MCI) and healthy controls (HC). We further investigated whether SDB was associated with advanced brain aging. We included a total of 330 participants, divided based on self-reported history of SDB, and matched across diagnoses for age, sex and presence of the ApoE4 allele, from the Alzheimer’s Disease Neuroimaging Initiative. Gray-matter volume was measured using voxel-wise morphometry and differences reflecting SDB, cognitive status, and their interaction were evaluated. Further, using an age-prediction model fitted on gray-matter data of external datasets, we predicted study participants’ age from their structural scans. Cognitive decline (MCI/AD diagnosis) and advanced age were associated with lower gray matter volume in various regions, particularly in the bilateral temporal lobes. BrainAGE was well predicted from the morphological data in HC and, as expected, elevated in MCI and particularly in AD. However, there was neither a significant difference between regional gray matter volume in any diagnostic group related to the SDB status nor an SDB-by-cognitive status interaction. Also, we found neither a significant difference in BrainAGE gap (estimated - chronological age) related to SDB nor an SDB-by-cognitive status interaction. In summary, contrary to our expectations, we were not able to find a general nor a diagnostic specific effect on either gray-matter volumetric or brain aging.

**Statement of significance:** Dementia syndromes including Alzheimer’s disease (AD), are a major global concern, and unraveling modifiable predisposing risk factors is indispensable. Sleep-disordered breathing (SDB) and its most prevalent form, obstructive sleep apnea, are suggested as modifiable risk factors of AD, but their contribution to AD hallmarks, like brain atrophy and advanced brain aging, is not clear to this day. While self-reported SDB is suggested to propagate aging process and cognitive decline to AD in clinical studies, here, we demonstrated that, SDB might not necessarily associate to brain atrophy and the advanced brain aging assessed by morphological data, in AD progession. However, multimodal longitudinal studies with polysomnographic assessment of SDB are needed to confirm such fundings.

## Introduction

Dementia syndromes including Alzheimer’s disease (AD), are a major global concern, with prevalence of 712 cases per 100,000 population in 2016, affecting 40-50 million people worldwide^1^. Considering that the numbers of AD patients have been more than doubled during past three decades^1^, it is critical to unravel the predisposing risk factors ^2^. These include advanced aging of the world population, but also modifiable risk factors such as cardiovascular disease, diabetes^2^, obesity^3^, and potentially sleep-disordered breathing (SDB)^4^. SDB ranges from partial (episodical) to complete airway obstruction leading to intermittent hypoxia, sleep fragmentation and intrathoracic pressure swings^5^. A bidirectional relationship has been proposed for SDB, including its most common form, obstructive sleep apnea (OSA), and AD. In particular, it has been suggested that patients with OSA are more likely to develop mild cognitive impairment (MCI) or dementia^6,7^. Moreover, a meta-analysis demonstrated that prevalence of OSA is five times higher in patients with AD than cognitively unimpaired individuals of the same age^4^.

Gray matter atrophy is a prime feature of pathologic brain aging^8^ and a well-known finding in AD, starting primarily in the medial temporal region and then globally affecting the brain as the disease progresses^9–11^. Morphometric analysis of the structural magnetic resonance images (MRI) has shown to reliably reveal this effect^12^. While some studies have shown gray matter atrophy in brain regions like the hippocampus, a key region involved in AD, to be associated with SDB in non-demented subjects^13–16^, others have shown either null results^17^ or paradoxical hypertrophy or thickening of gray matter in SDB^18–24^. Discrepancy between these findings is attributed to variations in cognitive status of participants, definitions of SDB severity, and method of gray matter volume assessment^14–16,18–24^. Thus, it remains unclear, whether SDB may result in brain atrophy similar to the volume changes in AD and hence contribute to its pathophysiology.

Aside from regional atrophy of the medial temporal lobe, AD is associated with advanced multivariate patterns of brain aging. In particular, it has been shown that individual subjects’ age can be predicted from gray matter morphometry in the cognitively normal population using machine-learning approaches^25^. That is, models trained to predict individuals ages based on larger cohorts of reference scans allow to estimate the age of a new person with a mean accuracy of 4-5 years^26^, while neurodegenerative disorders show a reliable pattern of advanced aging, i.e., a positive BrainAGE score (difference between age predicted based on the morphometric pattern and chronological age)^25,27–29^. Whether accelerated brain aging as seen in AD and to a lesser degree MCI is also present in potentially related conditions such as SDB is still an open question.

The aim of the current study is to shed further light on the potential relationship between brain atrophy patterns in SDB and AD at the regional and global level, answering two questions. 1) Do patients with SDB show grey matter atrophy across or in interaction with cognitive status (healthy control (HC), MCI, AD)? 2) Do patients with SDB show advanced brain aging across or in interaction with cognitive status (HC, MCI, AD)? To this end, we used data from the Alzheimer’s Disease Neuroimaging Initiative (ADNI), and established the validity of our methods by replicating previous findings for both aims in MCI and AD, and then assessed gray matter volume and BrainAGE differences between SBD+ and SBD-, including interactions with cognitive status.

## Methods

### Participants

Subjects were drawn from the Alzheimer’s Disease Neuroimaging Initiative (ADNI) database (adni.loni.usc.edu)^30^ based on their cognitive status and the medical history regarding SDB^7^. Diagnoses of MCI and AD were based on the ADNI criteria. Subjects with self-reported “sleep apnea” or “obstructive sleep apnea” or “OSA” symptoms or receiving treatment with “Continuous Positive Airway Pressure” (or “CPAP”) or “Bilevel Positive Airway Pressure” (or “BiPAP”/“BPAP”) were labeled as “SDB+”. wo independent physicians reviewed medical history to confirm diagnosis and grouping the subjects. Demographic and clinical variables were extracted for all individuals, missing covariate data were assessed and imputation was used for 5 participants with missing data-points. Using 1:1 propensity score matching method, we assembled 6 distinct sub-groups according to their cognitive (HC, MCI, AD) and SDB (SDB+ and SDB-) status. Covariates included in the matching were age, sex, years of education, body-mass index, cognitive status (AD/MCI/HC), presence of the Apolipoprotein E4 (ApoE4) allele, history of SDB treatment (only when matching between SDB+ subjects), T1 imaging protocol and, field strength (Table 1). Only subjects that passed the quality assessment tools of the CAT toolbox, including weighted image quality rating based on basic image properties and noise and geometric distortions, as well as checking homogeneity through the sample, were considered.

**Table 1.** Characteristics of the study subjects

|  | **SDB-** | **SDB+** | **P value** |
| --- | --- | --- | --- |
|  | N: 165 | n:165 |  |
| **Age (Mean (SD))** | 73.99 (7.70) | 74.91 (7.18) | 0.26 |
| **Sex, Female (%)** | 61 (37.0) | 48 (29.1) | 0.16 |
| **Cognitive status (%)** |  |  | 1.00 |
| **AD** | 24 (14.5) | 24 (14.5) |  |
| **MCI** | 111 (67.3) | 111 (67.3) |  |
| **HC** | 30 (18.2) | 30 (18.2) |  |
| **BMI (Mean (SD))** | 28.97 (5.95) | 29.08 (5.45) | 0.86 |
| **Education years (Mean (SD))** | 16.07 (2.75) | 16.16 (2.65) | 0.74 |
| **Handedness = Left (%)** | 18 (10.9) | 18 (10.9) | 1.00 |
| **Apoe4 allele count (%)** |  |  | 0.13 |
| **0** | 71 (46.7) | 94 (58.0) |  |
| **1** | 64 (42.1) | 53 (32.7) |  |
| **2** | 17 (11.2) | 15 (9.3) |  |
| **MMSE (Mean (SD))\*** | 26.07 (4.13) | 25.44 (4.93) | 0.25 |
| **CPAP/Surgery (%)\*** | 0 (0.0) | 56 (33.9) | – |
| **Protocol, MP-RAGE (%)** | 118 (71.5) | 124 (75.2) | 0.53 |

SDB: Sleep-disordered breathing, AD: Alzheimer's disease, MCI: Mild cognitive impairment, HC: healthy control, BMI: Body-mass index, MMSE: Mini-mental state examination, MP-RAGE: 3D magnetization prepared rapid gradient echo, CPAP: Continuous positive airway pressure
\*Not included in the matching

### Imaging acquisition and preprocessing

Participants had undergone a standardized protocol for high-resolution MRI T1 scans of the brain as previously described^31^. T1 imaging acquisition parameters were: TR= 2400 ms, minimum full TE, TI=1000 ms, flip angle= 8°, 24 cm field of view, acquisition matrix of 192 ×192 ×166 and with 1.25 ×1.25 ×1.2 mm3 slice size. We used Computational Anatomy Toolbox (CAT) v12^32^ and SPM12 (www.fil.ac.uk/spm) to perform voxel-based morphometry (VBM). This included correcting the bias-field distortions and noise removal, skull stripping, normalization to standard space and brain tissue segmentation into grey matter, white matter, and cerebrospinal fluid. Grey matter segments were modulated to represent actual gray matter volume. We then performed a biologically informed compression of the VBM data to 637 gray matter parcels based existing in-vivo brain parcellation^33^,^34^. Thus, grey matter volume of each participant was represented by 637 features each representing an individual parcel volume of that participant. All consecutive analyses were performed on this data.

### Statistical analysis of gray matter volume

Statistical analysis of gray matter volume of parcels included three consecutive parts, similar to approach used by Bludau et. al^35^; generating reference statistics, permuted statistics, and a family-wise error (FWE) correction for multiple comparisons. Here we used an n-way analysis of variance, to test effect of age, cognitive status (AD/MCI/HC), SDB status and SDB-by-cognitive status interaction, separately as independent variables (factors), on gray matter volume of each parcel as dependent variable. The F values (per parcel) of this ANOVA were considered as the reference statistics. In the subsequent permutation statistics for each factor, we randomly shuffled the labels for that factor 10,000 times, replicated the analysis and recorded the F-values to build a null-distribution. The comparison of the reference statistic with this distribution then allows non-parametric inference per parcel and factor, yielding uncorrected p-values. Importantly, however, we also recorded, per replication of the permutation, the highest statistics in the random data across the entire set, i.e., 637 brain regions, building a null-distribution for family-wise error correction. The threshold corresponding to p_FWE_ < 0.05 was then provided by the (set-wise maximum) value exceeded only in 5% of the replications.

### Age prediction

Brain age was estimated from the atlas-based representations of individual brain anatomy using a support vector machine (SVM) ensemble model. An independent (reference) large dataset consisting of 2089 (Figure 1A.1) subjects (between 55 and 85 years old) was compiled from several large public and private datasets including 1000Brains^36^, Cambridge Centre for Ageing and Neuroscience or Cam-CAN ^37^, OpenfMRI^38^, Dallas Lifespan Brain Study or DLBS, Consortium for Reliability and Reproducibility or CoRR^39^, IXI, and Enhanced Nathan Kline Institute-Rockland Sample or eNKI-RS^40^. Given the imbalance between age brackets, sites, and sex, we performed a stratified subsampling, choosing the same number of men and women, as well as similar numbers across age-brackets and a maximum of 30 subjects per age-bracket and sex per site. The actual subjects sampled in each replication from the overall database were drawn from the pool independently at random without replacement. Each of these sampled sets was then used to fit an individual SVM providing a weak learner for the ensemble which was applied to the test-data, i.e., the ADNI sample. The process was repeated 10,000 times, yielding 10,000 age predictions based on models trained on (different) balanced subsamples of the multicohort reference data. These predictions were then averaged (“bagging”) to yield the final age prediction based on the 637-parcel representation of the VBM data^41^. Each subjects BrainAGE score was finally calculated as bagged predicted age minus chronological age for each subject (Figure 1).

**Figure 1.**
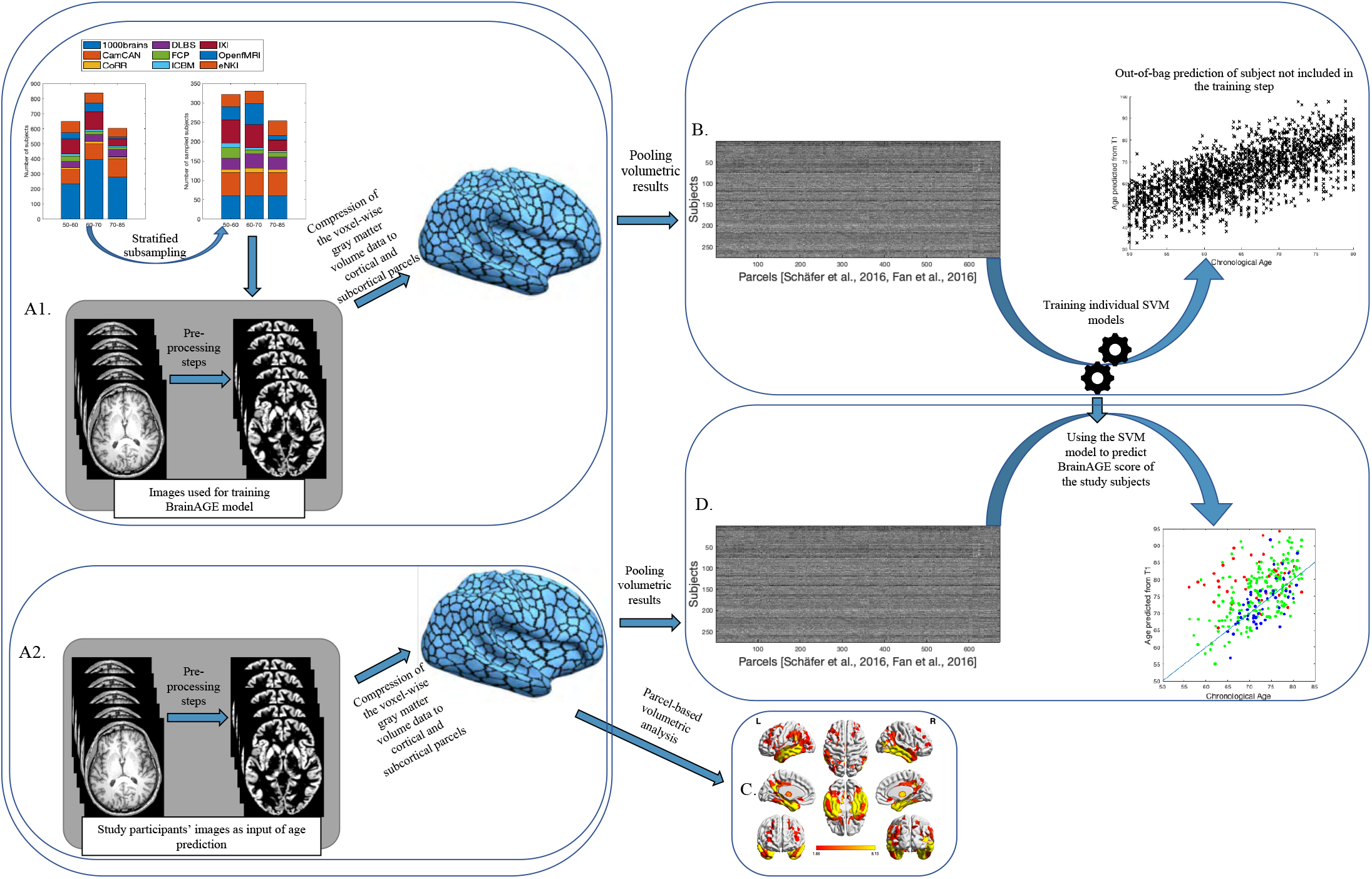
Main processing steps for parcel based volumetric study and age prediction based on gray matter morphometry. **A1**. T1 brain images of 2089 non-demented age, sex, and site stratified subjects were acquired through several imaging databases for development of age-prediction model (Training images). To obtain voxel-based gray matter volume data, standard pre-processing steps including normalization, segmentation and modulation for non-linear transformations have been done using Computational Anatomical Toolbox 12 (CAT12). A biologically informed compression of the voxel-wise gray matter volume data to 600 cortical and 37 subcortical regions was applied accordingly. **B.** Parcel-based results were then used as input for training the support vector machine (SVM) used for age-prediction model. **A2**. Similar pre-processing steps were done on T1 brain images of study-specific SDB+ and SDB-subjects (Study-specific images). Parcel-based results were used in two parallel analyses; 1) **C**. inputted to partial ANOVA tests for gray matter volume assessment according to presence of SDB and cognitive status as contrasts and 2) **D.** Decomposed with an OPNMF approach and inputted in the age prediction SVM model developed on the training images.

## Results

Both SDB+ and SDB-groups comprised of 24 AD, 111 MCI, and 30 HC participants, respectively. As enforced through the matching, there was no statistically significant difference in demographic variables, cognitive status, and presence of the ApoE4 allele between SDB groups. Table 1 summarizes the characteristics of all study groups.

### Effects on grey matter volume

As noted in the methods, association of parcel-wise gray matter volume with age, cognitive status, SDB status, and SDB-by-cognitive status interaction was assessed using non-parametric inference with FWE correction for multiple comparisons. There were strong (P_FWE_ <0.001) and widespread negative associations of regional grey matter volume with age, in particular in the bilateral temporal lobes, bilateral prefrontal, middle and superior frontal areas, bilateral medial and lateral occipital areas, cerebellum and thalamus, caudate and putamen in the subcortical gray matter (Figure 2A). The cognitive status was significantly associated with reduced gray matter volume in many bilateral parcels with dominancy in the left hemisphere (P_FWE_<0.001). Bilateral temporal lobes including fusiform gyri, medial temporal lobes and hippocampal formations, and inferior and middle temporal lobes had significantly lower volume in participants with MCI and particularly AD. Moreover, reduced gray matter volume was seen in bilateral insula, middle frontal, and cingulate cortices, as well as left superior frontal cortex (Figure 2B). In turn, when testing for effects of SDB status and SDB-by-cognitive status interaction, we found no significant regions anywhere in the brain (all P_FWE_ > 0.05).

**Figure 2.**
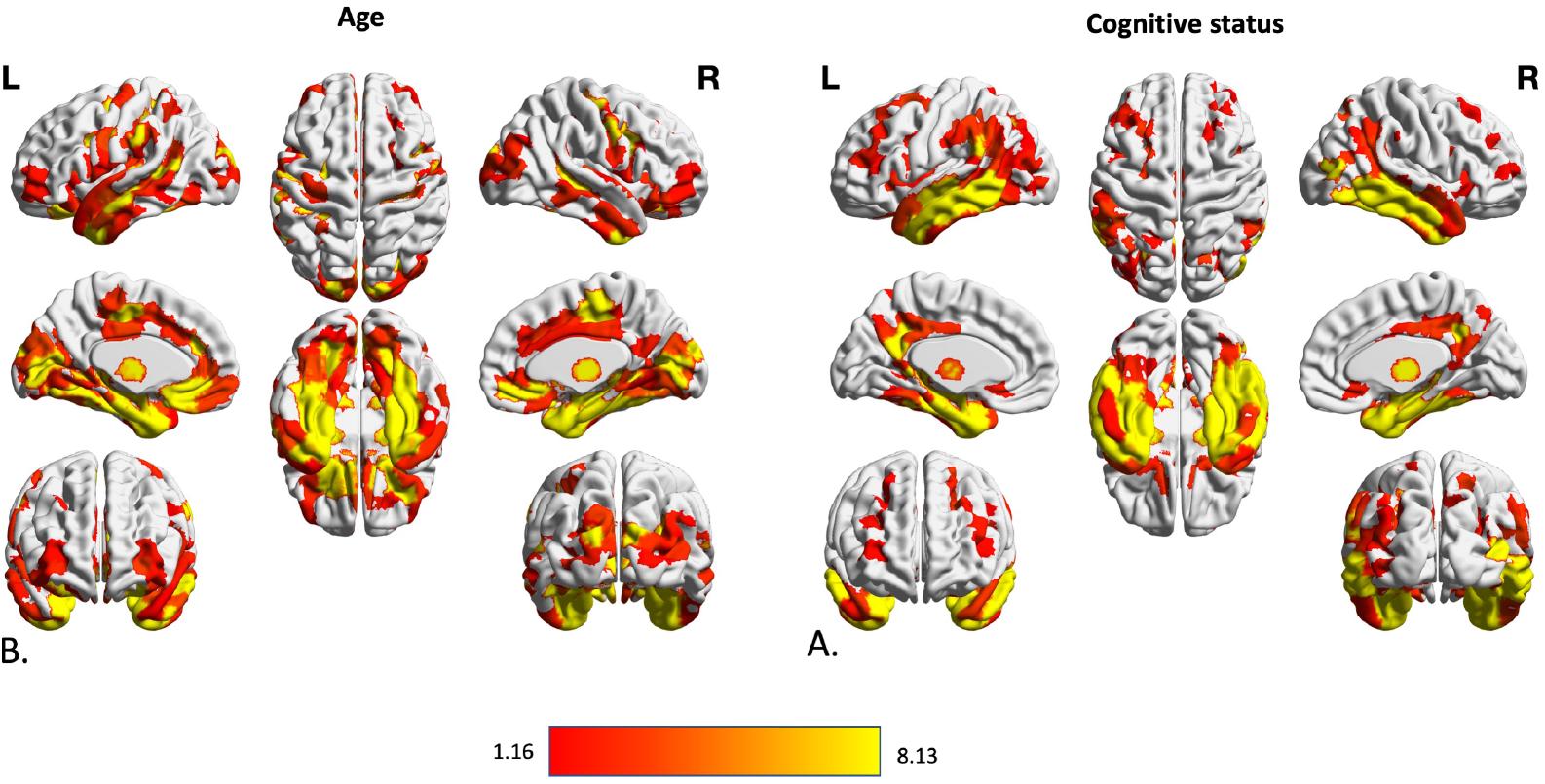
Association between volumetric data of cortical and subcortical parcels and age and cognitive status of subjects. Gray matter volume differences in 600 cortical parcels and 37 subcortical volume was assessed using three steps of using F value of an n-way analysis of variance as reference statistics, running 10,000 permutations per randomly shuffling different parcels, under assumption of label exchangeability, and correction of p values using family wise error (FWE) method. Significant parcels are illustrated as the heated areas on the brain maps considering (A) age and (B) cognitive status. Since there were no significant results regarding SDB presence or SDB-by-diagnosis interaction, results according these factors have not been illustrated here.

### Effects on estimated brain age

The mean absolute error between predicted and chronological age in the HC group was 3.59 years, indicative of the very good performance of the ensemble prediction model. We then calculated the BrainAGE score as the per-subject difference between predicted and chronological age and tested for its association with cognitive status, SDB status, and the SDB-by-cognitive status interaction. As it is shown in Figure 3, participants with MCI and in particular AD showed an advanced brain age (on average 4.0 and 9.1 years, respectively), in line with previous studies. However, there was no significant effect on BrainAGE scores associated with SDB status, nor was there a positive SDB-by-cognitive status interaction suggesting that SDB may not lead to advanced brain aging (Figure 3C).

**Figure 3.**
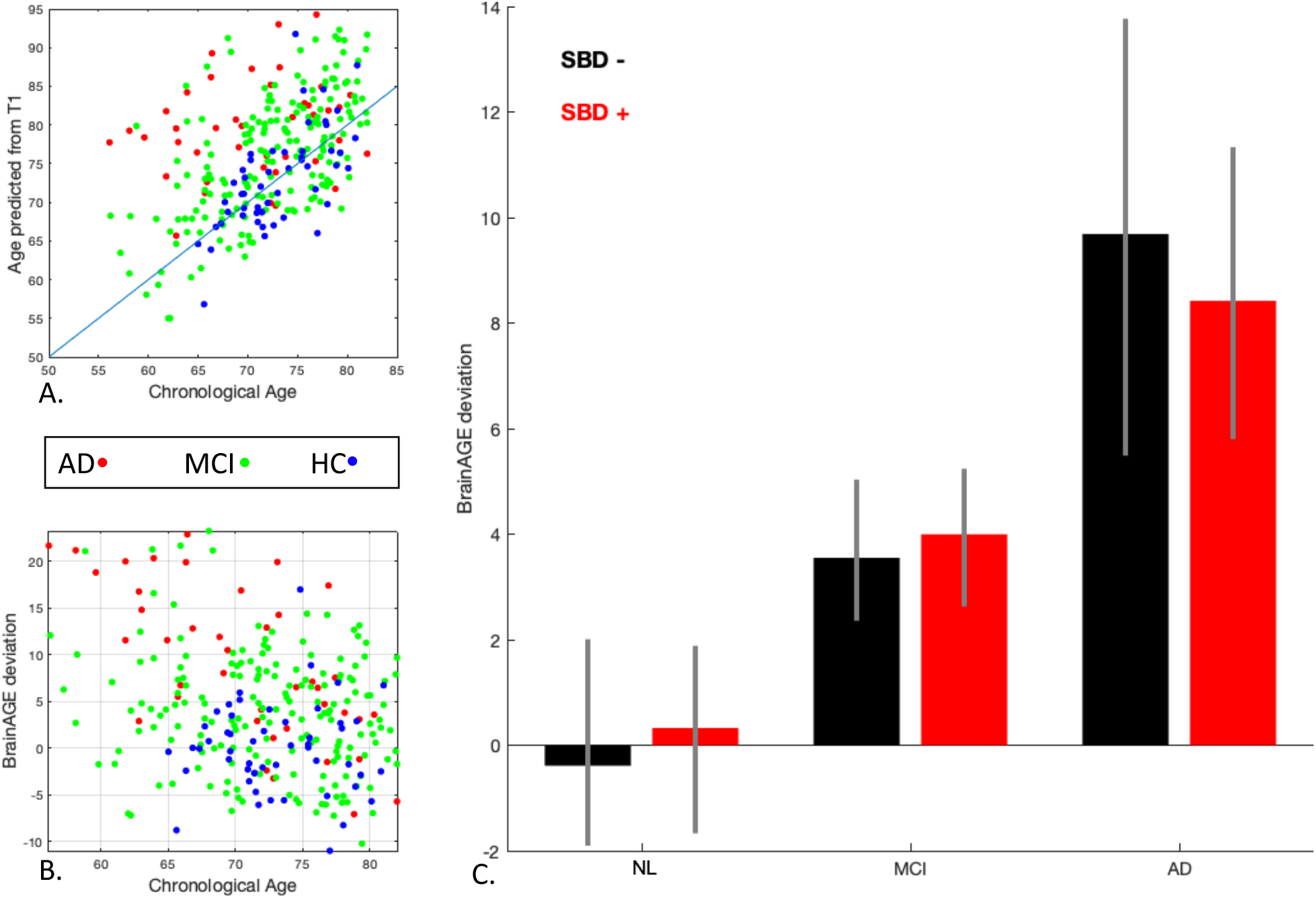
Results of the BrainAGE prediction method based on the presence of SDB and cognitive status. **A.** Relationship between chronological age and the predicted age from T1 images in the AD, MCI and HC groups. There is an evident higher predicted age for the participants AD and MCI compared to HC group, in accordance with advanced pathological brain aging in the AD course. **B.** The BrainAGE score shows positive and bigger deviation from chronological age in the AD and MCI groups. **C.** Despite the significantly higher BrainAGE deviation associated with AD and MCI, no significant deviation was seen between BrainAGE score of SDB subgroups.

## Discussion

Our findings confirmed previously reported gray matter atrophy and accelerated biological brain aging in patients with MCI and AD, corroborating the robustness and validity of our analytical approach. Importantly, we were not able to demonstrate any effect of SDB, independently or in interaction with cognitive status, on either regional grey matter volume or brain aging score. Several limitations however, may compromise the interpretation of our results. Sample sizes of SDB+ subjects in the HC and AD groups were small. Moreover, the groups were heterogeneous in terms of clinical characteristics and imaging specifications. We used propensity-score matching and stratified subsampling of external datasets to minimize the effects of heterogeneity. As previously mentioned on publications using the ADNI database^7,42^, the self-reported measure of SDB can be influenced by both the recall bias of cognitively impaired subjects as well as by a high prevalence of undiagnosed OSA in the general population, therefore increasing the probability of false negative cases in the SDB-groups^7^. Moreover, assessment of the severity of SDB and disease duration were not available.

### Grey matter volume alterations in AD and SDB

One of the main characteristics of MCI and AD is generalized gray matter loss in the brain, which mostly starts in the medial temporal lobe and multimodal association areas^8–10^. Neuroimaging meta-analyses have demonstrated atrophy in the medial temporal lobe, limbic regions (left parahippocampl gyrus, left posterior cingulate gyrus, amygdala and uncus), thalamus, temporal, parietal, frontal and cingulate cortices^43,44^. A similar but milder distribution of gray matter atrophy is evident in brain of patients with MCI^43,45^. In accordance with the previous brain volumetric studies, we found diffuse gray matter loss in MCI and AD. The atrophy was mainly located in the bilateral temporal lobe and medial temporal areas with higher intensity in AD compared to MCI.

Assessing the volumetric changes due to SDB, we did not observe any significant alteration in gray matter volume, neither in HC subjects, nor in patients with MCI or AD. Furthermore, self-reported SDB interaction with cognitive status (i.e. HC, MCI or AD) showed no associations with gray matter volume. Historically, there has been an inability to replicate results among the brain imaging studies of SDB in non-demented populations. While several studies have reported gray matter atrophy in the insula, amygdala, middle and lateral temporal regions, and cerebellum in non-demented populations with SDB^13–16,46–48^, others have either shown no associations^17,49^ or paradoxical enhancement in the gray matter volume of regions like the motor cortices, prefrontal cortex, thalamus, putamen, and the hippocampus^20–24,47^. In addition, there is a general lack of longitudinal studies, which would enable the study of non-linear associations between SDB and cortical atrophy as suggested by these cross-sectional findings. Despite these important gaps in the literature, three meta-analyses have concluded that OSA is associated with gray matter atrophy in a few selected regions including the cingulate, amygdala, hippocampus, right central insula, right middle temporal gyrus, right premotor cortex, and cerebellum^13,50,51^.

The observed null association between SDB and gray matter volume should however be interpreted with caution. First, it has been suggested that aging may have partially protective mechanisms against SDB, such as reduced production of oxidative stress after apneas and decreased blood pressure and heart rate responses after arousals^23^. The average old age of ADNI subjects (~75 years-old) could therefore explain this non-significant association between SDB and brain morphometry. Despite numerous studies and meta-analyses focused on the changes in gray matter in middle-aged patients with OSA, there are few studies on gray matter changes in older adults with SDB and neither have found any decreases in thickness or volume in cortical gray matter^52–54^. Second, it is possible that SDB-related brain damage impacts more selectively brain function^55^ or amyloid burden^17^ than gray matter volume alone, or that differential diagnosis between *SDB-related* and *age-related* brain atrophy is difficult in single-point observational studies, particulary in those cases in which groups are matched by age and cognitive status. Third, this could also be a sign of: a) survival bias, as most SDB+ may have transitioned to AD and only those with very low cortical atrophy or high in cognitive reserve at disease onset would remain as HC or MCI at cross-section; or, b) selection bias due to matching by the ApoE4 allele, as it has been reported that the ApoE4 allele interacts with brain aging scores measured by the BrainAGE method, revealing potential neuronal compensation in healthy ApoE4+ adults^73^, which could also result in null findings. Fourth, we did not account for other comorbidities and possible confounders alongside age or presence of the ApoE4 allele in the prediction models^72^. Finally, previous MRI studies mostly recruited patients with PSG-diagnosed OSA from sleep clinics, which might be a different population from those recruited in memory clinics with selfreported assessment of SDB based on clinical interview.

We were also not able to demonstrate any interaction between SDB and MCI or AD with brain atrophy. This is indicative that despite the frequent clinical co-occurrence of SDB and AD, there may be no synergy between them in accelerating gray matter atrophy. Recent investigations using cerebrospinal fluid and PET imaging suggest an interplay between amyloid production/clearance and SDB^17,42,56–58^. These include an impairment in the cerebrospinal fluid-interstitial fluid exchange^60^, cerebral edema secondary to an intermittent hypoxia^61^ (similar to the increase in brain volume and pseudoatrophy observed in multiple sclerosis), and compensatory excessive neuronal synaptic activity^62^ in SDB, all of which could potentially lead to an increase in beta-amyloid deposition and its clearance reduction. It is therefore possible that the presence of SDB is associated with AD risk only through beta-amyloid deposition^42,58^ or altered brain function^63–65^, but an interaction should have been observed in MCI or AD where it is generally accepted that neuronal loss follows amyloid deposition. More studies are needed to better understand the compensatory increase in gray matter volume in SDB suggested by several studies, as well as the precise progression of brain atrophy in AD, as both may have contributed to obtaining such negative findings.

### BrainAGE prediction in AD and SDB

Brain age prediction methods have been used in cognitively normal subjects^26,66^ and several studies have used the ADNI dataset with mean absolute error (MAE) ranging from 3 to 6 years^25,27^. We implemented an advanced sensitive BrainAGE estimation method to detect pathologic brain aging, using repeated SVM models fitted on parcel-wise gray matter volume data of on stratified subsamples from external cohorts, making the model less sensitive to heterogeneity in images^25^. Compared to previous studies, while using multiple datasets for training prediction model, our age prediction results were accurate with an MAE of 3.6 years in HCs. Replication of previous findings in AD, taken together with acceptable MAE, is indicative of reliability of our proposed method in gray matter volume assessment and age estimation

While there is no exact definition for accelerated brain aging, BrainAGE score has been shown to be a sensitive predictor of disease progression in dementia^27–29^. Previous findings on increased BrainAGE score in MCI and AD course^67–69^, are in agreement with the reported accelerated aging of the demented brain shown in-vivo and ex-vivo studies^70^. The BrainAGE score in studies using ADNI ranged from almost zero for patients with stable MCI, to 5.7–6.2 years for patients with progressive MCI, and reached up to 10 years for patients with AD^27^. We found the average 4.1 and 9 BrainAGE scores in patients with AD and MCI, in agreement to previous findings using ADNI data. Since we did not distinguish patients with progressive from stable MCI, our results in the MCI group were modest compared to other studies including patients with late or progressive MCI.

## Conclusions

In summary, we have confirmed the acceleration of brain atrophy and advanced brain aging in MCI and AD participants from the ADNI cohort compared to healthy controls. We further found that self-reported SDB in subjects with a diagnosis of HC, MCI or AD was neither associated with gray matter volume reduction, nor with accelerated brain aging. While SDB is suggested to propagate the aging process, amyloid burden and cognitive decline to AD, it may not necessarily associate to brain atrophy and the estimated brain age in AD progession.

## Acknowledgment

The authors gratefully acknowledge the efforts, time, and dedication of the participants and staff of the Alzheimer’s Disease Neuroimaging Initiative (ADNI). Collection and sharing of data for this project was funded by ADNI (National Institutes of Health Grant U01 AG024904) and DOD ADNI (Department of Defense award number W81XWH-12-20012). ADNI is funded by the National Institute on Aging, the National Institute of Biomedical Imaging and Bioengineering, and through generous contributions from the following: AbbVie, Alzheimer’s Association; Alzheimer’s Drug Discovery Foundation; Araclon Biotech; BioClinica, Inc.; Biogen; Bristol-Myers Squibb Company; CereSpir, Inc.; Cogstate; Eisai Inc.; Elan Pharmaceuticals, Inc.; Eli Lilly and Company; EuroImmun; F. Hoffmann-La Roche Ltd and its affiliated company Genentech, Inc.; Fujirebio; GE Healthcare; IXICO Ltd.; Janssen Alzheimer Immunotherapy Research & Development, LLC.; Johnson & Johnson Pharmaceutical Research & Development LLC.; Lumosity; Lundbeck; Merck & Co., Inc.; Meso Scale Diagnostics, LLC.; NeuroRx Research; Neurotrack Technologies; Novartis Pharmaceuticals Corporation; Pfizer Inc.; Piramal Imaging; Servier; Takeda Pharmaceutical Company; and Transition Therapeutics. The Canadian Institutes of Health Research is providing funds to support ADNI clinical sites in Canada. Private sector contributions are facilitated by the Foundation for the National Institutes of Health (www.fnih.org). The grantee organization is the Northern California Institute for Research and Education, and the study is coordinated by the Alzheimer’s Therapeutic Research Institute at the University of Southern California. ADNI data are disseminated by the Laboratory for Neuro Imaging at the University of Southern California. Of note, Simon B. Eickhoff is supported by the Deutsche Forschungsgemeinschaft, the National Institute of Mental Health (R01-MH074457), the Helmholtz Portfolio Theme “Supercomputing and Modeling for the Human Brain” and the European Union’s Horizon 2020 Research and Innovation Programme under Grant Agreement No. 7202070 (HBP SGA1). Ivana Rosenzweig was supported by the Wellcome Trust [103952/Z/14/Z]. Ricardo Osorio’s salary is supported by NIH/NIA’a R01AG056031, R01AG056531 and R21AG055002.

## Abbreviations

MRI: Magnetic resonance imaging
SDB: Sleep-disordered breathing
OSA: Obstructive Sleep Apnea
AD: Alzheimer’s disease
MCI: Mild cognitive impairment
HC: Healthy Control
ApoE4: Apolipoprotein E4
MMSE: Mini-Mental State Examination
SVM: Support vector machine
VBM: Voxel-based morphometry
CAT: Computational Anatomy Toolbox
SPM: Statistical Parametric Mapping
FWE: Family-wise error
ANOVA: Analysis of variance

